# Circadian control in the timing of critical periods during *Drosophila* larval neuronal development

**DOI:** 10.1101/2024.03.21.586088

**Authors:** Sarah Doran, Adam A. Bradlaugh, Jack Corke, Richard A. Baines

**Affiliations:** Division of Neuroscience, School of Biological Sciences, Faculty of Biology, Medicine and Health, University of Manchester, Manchester Academic Health Science Centre, Manchester, M13 9PL, UK

**Keywords:** Cryptochrome, PDF, critical period, circadian clock

## Abstract

Critical periods (CPs) of development are temporal windows of heightened neural plasticity. Activity perturbation during CPs can produce significant, and permanent, alterations to the development of neural circuits. In this study we report a circadian mechanism underlying the timing of CPs in *Drosophila* embryonic and larval development. These CPs occur at ∼24 hr intervals and are open to manipulation through blue light (BL)-activation of the circadian regulator Cryptochrome (CRY). This manipulation is sufficient to destabilize the larval CNS, evidenced by an induced seizure phenotype when tested at third instar (L3). In addition to CRY nulls, genetic ablation of the *period* gene also mitigates the BL exposure seizure phenotype and, moreover, alleles of *period* that affect circadian timing alter the timing of the CPs. Our analysis shows a clear role for the main clock neuropeptide, pigment dispersing factor (PDF), to transduce the output of these CPs. Targeted PDF receptor knockdown, in either GABAergic or CRY-positive neurons, is sufficient to prevent the CRY-mediated seizure phenotype. This study is a first demonstration of a circadian mechanism in *Drosophila* larvae, and whilst this alone is of major significance, our results highlight the potential of using *Drosophila* larvae as a model to investigate the impact of circadian rhythms on early neuronal development in higher organisms, which remains experimentally challenging.

**Significance Statement:** Whilst the role of the biological clock is well understood in adult organisms, the same is not true for embryonic development. How the maternal clock impacts the mammalian fetus remains poorly understood. Given that many expectant mothers experience altered circadian rhythms, largely due to nightshift working, it is important to address these concerns. Here we identify clock-mediated periods in neural development of the embryonic Drosophila which can be manipulated by light. These findings provide an experimental opportunity to better understand the role of the circadian clock in early development.

## Introduction

Developing neural circuits are maximally open to modification, including by external factors, during temporally defined periods. Such periods have been termed ‘critical’ or ‘sensitive’ based on their perceived influence. Manipulations during critical periods (CPs) often drive permanent change that becomes locked in. Similar manipulations during sensitive periods (SPs) can be equally profound but, differentially, can be modified following period closure (1–4). First conceptualized in the 1960s (5), CPs are developmental windows during which changes in neural activity support both anatomical and functional tuning of neural networks. This includes the possible encoding of homeostatic “set points” that ensure appropriate activity levels in both neurons and neural networks (1, 4, 6–8). Previous work from our group has identified a CP in the *Drosophila* embryo, during 17-19 hours after egg laying (AEL), during which activity-manipulation is sufficient to permanently change the stability of the locomotor network (6, 9). This change in stability is measured through the induction of a seizure phenotype, indicative of significant change to the larval CNS. Significantly, exposure to BL during this embryonic CP is similarly able to lead to an increased seizure phenotype, an effect that requires the presence of CRY (10).

SPs differ from CPs in that while experiences during these timeframes still can have significant influence on development and behavior, they are not so tightly bound by a specific timeframe (11). SPs are, thus, believed to be required to facilitate learning at key developmental milestones but, importantly, such learning can nevertheless still occur at other times. Pertinent examples of SPs include early imprinting and acquisition of language or motor skills in developing animals (2, 12). In human, these periods include childhood adversity during SPs predicting changes in DNA methylation (13), and growing evidence that there is a SP for emotional development in children (14). However, the differentiation and contribution to neural development of both types of plasticity-windows remains poorly understood.

We report that embryonic and larval development in *Drosophila* is punctuated by CPs at roughly 24 hr intervals, indicative of orchestration by the molecular clock. Circadian rhythms are integral to all living things and serve to increase an organism’s fitness. The prime purpose of the circadian clock is to coordinate activity and metabolism to anticipated changes in the daily light:dark (LD) cycle, using environmental factors such as light and/or temperature as zeitgebers (15). The molecular clock in *Drosophila* is a negative feedback loop in which transcription factors drive expression of their own repressors. This core transcription-translation feedback loop is comprised of four key elements: CLOCK (CLK), CYCLE (CYC), PERIOD (PER) and TIMLESS (TIM). CLK and CYC form a heterodimeric complex, which in turn promotes the transcription of the repressors *per* and *tim* (16, 17). Alongside this negative feedback loop ,CRY, a BL sensitive flavoprotein, entrains this molecular oscillation to the environmental LD cycle through light-mediated degradation of TIM (18). The time taken for the transcription-translation feedback loop to complete a full cycle is the foundation underlying the clock’s ability to keep time.

In this study, we identify a series of CPs, occurring at ∼24 hr intervals, beginning in embryonic through to larval development. These CPs are open to manipulation by BL and are absent in a CRY loss-of-function mutant. Moreover, genetic manipulation of *per*, or the principal clock signaling neuropeptide - PDF - uncovers a novel role for the circadian clock during the larval stages of *Drosophila* neural development.

## Results

### Blue light sensitive windows occur at 24-hour intervals through larval development

Wildtype Canton-S (CS) embryos were exposed to BL for 2 hr periods prior to, during, and after the characterized embryonic CP at 17-19 hrs AEL (4). Only exposure during 17-19 hrs was sufficient to induce a seizure phenotype at wandering L3 (one-way ANOVA, *f*(10, 224) = *1.213*, *p < 0.0001, n = 15*, Fig. 1A-B). These results validate our previous findings by showing that BL exposure during 17-19 hrs AEL is sufficient to induce a seizure phenotype when compared to dark reared controls (embryos that did not experience BL exposure during these hours) (6, 19). The seizure phenotype is indicative of an incorrectly tuned locomotor network and provides a reliable, and relatively simple assay to monitor for effects of activity-manipulation (6, 8, 19). To investigate whether the 17-19 hr window for BL sensitivity is a single occurrence within development, we exposed CS L1 larvae to BL for 6 hr intervals and measured their seizure phenotype at L3 (Fig. 1B). Where we predicted that a CP might occur (based on 24 hr circadian rhythmicity) we exposed larvae to BL for shorter 2 hr intervals. We found that L1 larvae exposed to BL at 20-22 hrs after larval hatching (ALH) showed an increased L3 seizure phenotype compared to dark reared controls (larvae not exposed to BL during this period, *p < 0.0001*). Exposure during L1 was similarly confined to this single 2hr period. Thus, L1 larvae exposed to BL at 18-20 hrs ALH (*p > 0.99*) or 22-24 hrs ALH (*p > 0.99*) did not exhibit a seizure phenotype. With the assumption that BL-sensitive CPs are 2 hrs long, we exposed larvae to BL, for 2 hr periods, at 24 hrs intervals after the effective L1 exposure time: thus 44-46 hrs and again at 68-70 hrs ALH. BL exposure at both time intervals increased seizure recovery time compared to dark reared controls (*p < 0.0001*). However, BL exposure either before, or after, each period (42-44 hrs and 46-48 hrs; 66-68 hrs and 70-72 hrs) did not (Fig 1B, Table 1). These results clearly show that there are windows of heightened neural plasticity, sensitive to BL manipulation, at ∼24-hour intervals throughout larval development. Moreover, exposure to BL during these windows is sufficient to destabilize the larval nervous system as evidenced by an induced seizure phenotype at wandering L3.

**Figure 1.**
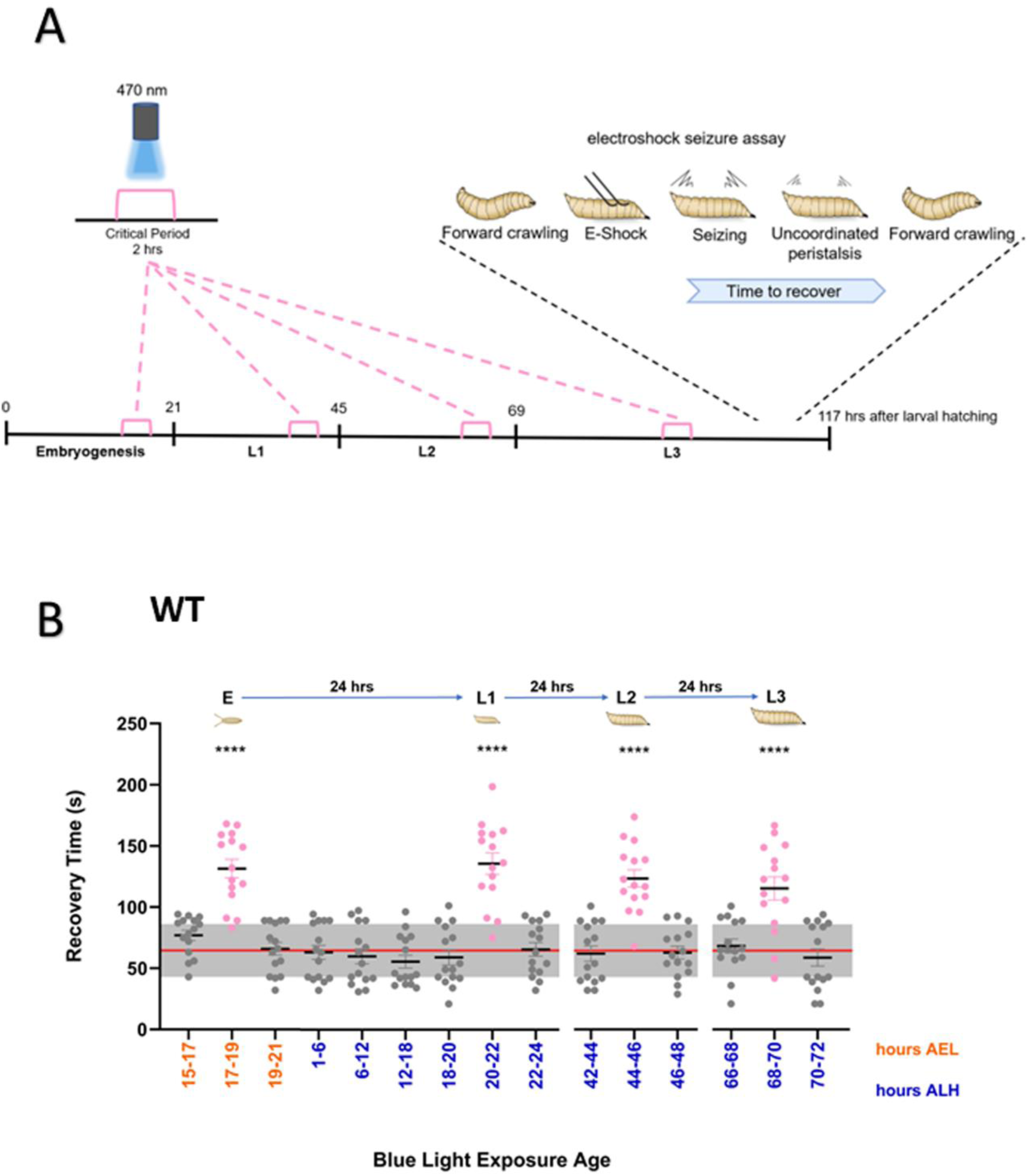
Blue light sensitive CPs occur at 24-hour intervals. (**A**) Schematic diagram of the electroshock protocol following BL exposure. Pink brackets identify the 2-hour time windows during which BL exposure occurred. Blue light exposure did not occur sequentially but separately during different experiments. Wandering L3 were challenged by electroshock to evoke a seizure-like phenotype. The recovery time for normal crawling behavior to resume was measured. (**B**) Seizure phenotypes were observed when BL exposure occurred at ∼24-hour intervals throughout development. CS embryos were exposed to BL at 2-hour intervals where a sensitive window was expected or at 6-hour intervals during L1 when not predicted. CS embryos exposed to BL at 17-19 hrs AEL exhibited a seizure phenotype (one-way ANOVA, *f(10,224)* = 1.213, *p < 0.0001, n = 15*, dark reared = red line, dark reared SEM = grey box). CS late-stage L1 larvae exposed to BL at 20-22 hrs ALH (*p < 0.0001, n = 15*), and L2 larvae exposed to BL for 2 hrs at 44-46 hrs ALH (*p < 0.0001, n = 15*), and 24 hrs after again for mid-stage L3 larvae exposed to BL at 68-70 hrs ALH (*p < 0.0001, n = 15)* all show an induced seizure phenotype. The grey data points depict the BL exposure groups that did not have an increased seizure recovery time at wandering L3 (full descriptive statistics and p values are reported in Table 1).

**Figure 2.**
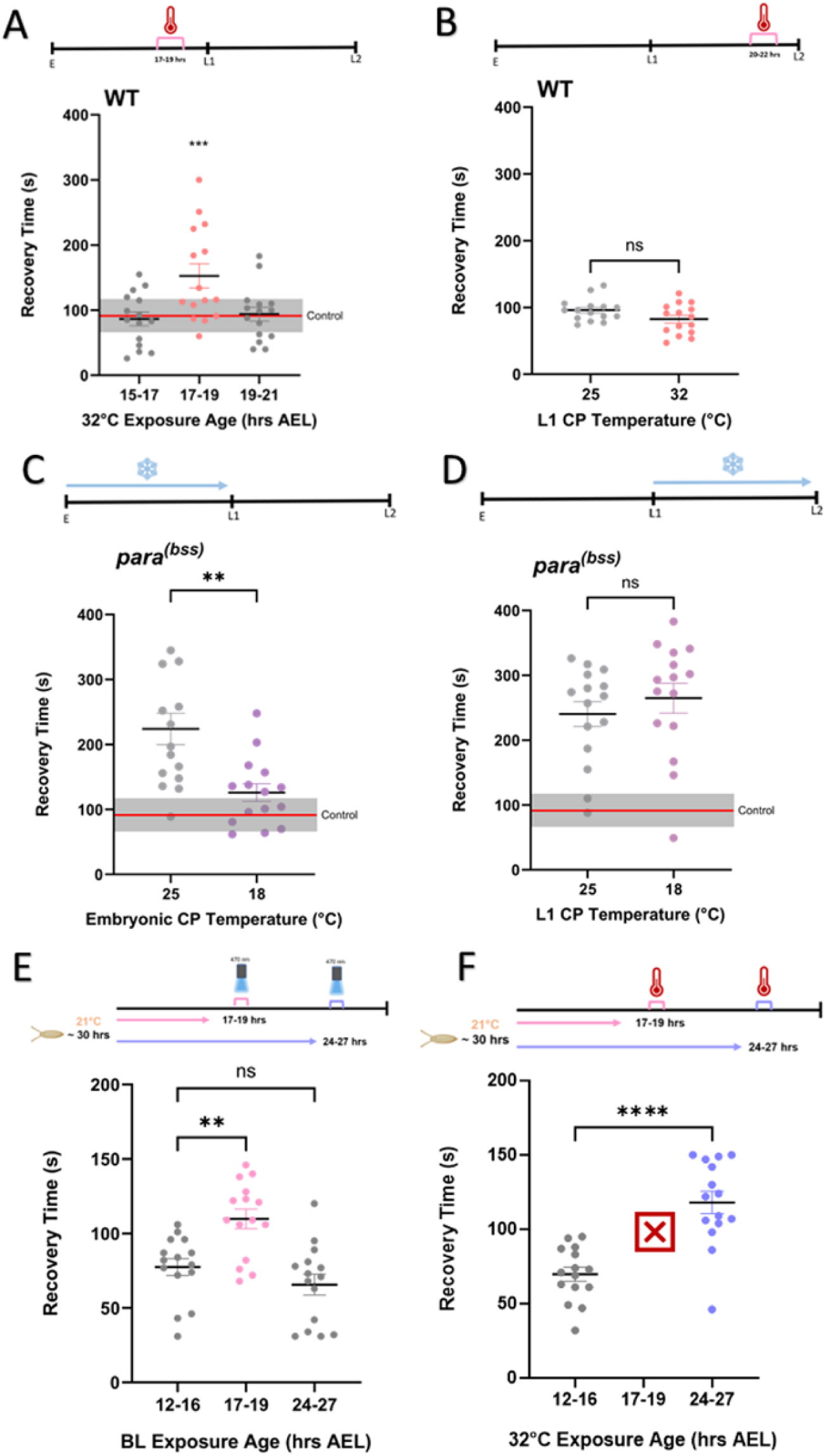
BL-sensitive CPs differ to a previously characterized embryonic activity-CP. (A) CS embryos exposed to 32°C, between 17-19 hrs AEL, show a higher seizure recovery time when electroshocked at wandering L3 (One-way ANOVA f(3, 56) = 4.715, p = 0.0001, n = 15). Exposure to 32°C at 15-17 hrs AEL (p = 0.858) or 19-21 hrs AEL (p = 0.575) was without effect. Control range (± sem) are dark-reared (gray horizontal bar). (B) CS late-stage L1 larvae exposed to 32°C at 20-22 hrs ALH did not show an increased recovery time (unpaired t-test, t(28) = 1.88, p = 0.071, n = 15). (C) parabss, reared at 18°C between 17-19 hrs AEL, show a significantly reduced seizure recovery time compared to a 25°C reared control (unpaired t-test, t(28) = 3.523, p = 0.0015, n = 15). (D). parabss L1 larvae, reared at 18°C between 20-22 hrs ALH, did not show a lower recovery time than a 25°C reared control (unpaired t-test, t(28) = 0.817, p = 0.421, n = 15). (E) The embryonic BL-CP is time locked at 17-19 hrs, regardless of developmental temperature. There was an increase in recovery time in 21°C reared embryos exposed to BL at 17-19 hrs AEL (one-way ANOVA, f(2, 42) = 0.462, p = 0.0017, n = 15) but not at 24-27 hrs AEL (p = 0.333). (F) The embryonic activity-CP is developmentally locked. The increase in seizure recovery time, due to 32°C exposure, occurred later at 24-27 hrs AEL in embryos developing at 21°C (unpaired t-test, t(28) = 5.45, p < 0.0001, n = 15). 32°C exposure between 17-19 hrs AEL was lethal.

**Table 1.** Full Descriptive Statistics of Figure 1B.

| BL exposure age (AEL/ALH) | Mean recovery time (s) | P value | SEM | Significance |
| --- | --- | --- | --- | --- |
| 15-17 | 76.87 | 0.310 | 4.149 | NS |
| 17-19 | 131.3 | < 0.0001 | 7.591 | **** |
| 19-21 | 65.93 | 0.992 | 5.218 | NS |
| **1-6** | 63 | > 0.9999 | 5.765 | NS |
| **6-12** | 59.80 | > 0.9999 | 6.082 | NS |
| **12-18** | 55.53 | > 0.9999 | 5.306 | NS |
| **18-20** | 58.93 | > 0.9999 | 6.214 | NS |
| **20-22** | 135.3 | < 0.0001 | 8.734 | **** |
| **22-24** | 65.47 | 0.9954 | 5.514 | NS |
| **42-44** | 62.13 | > 0.9999 | 6.256 | NS |
| **44-46** | 123.6 | < 0.0001 | 7.125 | **** |
| **46-48** | 62.93 | > 0.9999 | 5.200 | NS |
| **66-68** | 68.40 | 0.9313 | 5.645 | NS |
| **68-70** | 115.5 | < 0.0001 | 9.527 | **** |
| **70-72** | 58.80 | > 0.9999 | 7.016 | NS |
| **CONTROL** | 57.93 |  | 6.771 |  |

### BL-sensitive windows differ from the embryonic CP

The discovery of larval BL-sensitive CPs was unexpected. For clarity, we hereafter term these CPs as ‘BL-CPs’ to distinguish them from the previously characterized embryonic activity-dependent CP (hereafter termed ‘activity-CP’). An immediate question is whether BL-CPs are equivalent to the well-characterized embryonic activity-CP. In addition to BL, the embryonic activity-CP can be manipulated by genetics or pharmacology (6, 8). However, neither approach easily lends itself to temporal control. It has, however, recently been shown that the embryonic activity-CP can be manipulated by elevating ambient temperature (to 32°C, per comm. Matthias Landgraf, Cambridge). We similarly find that exposing CS embryos to 32°C, during 17-19 hrs AEL, is sufficient to induce a seizure phenotype at L3 (one-way ANOVA *f*(3, 56) = *4.76*, *p = 0.0001, n = 15*. Fig 2A). However, identical 32°C exposure is not sufficient to produce a seizure phenotype at 15-17 hrs AEL (*p = 0.86*) or at 19-21 hrs AEL (*p = 0.57*). Surprisingly, CS larvae exposed to 32°C, at late stage L1 during the BL-CP of 20-22 ALH, did not exhibit a seizure phenotype at L3 (unpaired t-test, *t*(28) = *1.88*, *p = 0.071, n = 15*, Fig. 2B). This indicates that the BL-CPs cannot be activity-manipulated in the same manner as the embryonic activity-CP.

To further investigate whether BL-CPs identified above are equivalent to the embryonic activity-CP, we utilized a characterized seizure mutant, para*^bangsenseless^ (para^bss^),* which we have previously shown to lose its characteristic seizure phenotype when reared at lower temperatures during the embryonic activity-CP (6). We validated this observation by showing that *para^bss^* embryos reared at 18°C, between 17-19 hrs AEL, have a decreased seizure recovery time at L3, compared to 25°C reared controls (unpaired t-test, *t*(28) = *3.52*, *p = 0.0015, n = 15*, Fig 2C). To investigate whether this same seizure rescue occurs during a larval BL-CP, *para^bss^* larvae were reared at 18°C between 18-24 hrs ALH (spanning the L1 BL-CP). We found that the *para^bss^* larvae reared at 18°C did not show a decreased recovery time (unpaired t-test, *t*(28) = *0.82*, *p = 0.421, n = 15*, Fig. 2D). This demonstrates, again, that larval BL-CPs differ from the previously defined embryonic activity-CP.

The opening of a canonical CP is dictated by a given neural network reaching a key developmental milestone, often coinciding with the maturation of inhibitory GABAergic interneurons (4, 20–22). In this instance, the embryonic activity-CP and BL-CP occur simultaneously between 17-19hrs AEL. Thus, we asked whether the activity-CP was locked to a specific developmental checkpoint, whilst we predicted that the BL-CP was temperature compensated as predicted for the circadian clock (23, 24). Embryogenesis lasts ∼21 hrs at 25°C, but is extended to ∼30 hrs at 21°C (6). We first exposed 21°C reared embryos to BL at either 17-19 hrs AEL or the developmentally adjusted time of 24-27 hrs AEL. We found that the timing of BL sensitivity did not change, remaining at the earlier 17-19 hr period (one-way ANOVA, *f*(2, 42) = *0.46*, *p = 0.0017, n = 15,* Fig 2E). BL-exposure between 24-27 hrs AEL was without effect (*p = 0.33*). This suggests that, despite the changes in temperature causing development to lengthen, the BL-CP is temperature compensated.

If, on the other hand, the embryonic activity-CP occurs at a specific developmental checkpoint, then its timing should shift accordingly from 17-19 hrs to 24-27 hrs AEL to compensate for the elongated development time. Intriguingly, exposure to 32°C between 17-19 hrs AEL, when development was delayed, was embryonic lethal. However, higher temperature exposure (32°C) between 24-27 hrs AEL was sufficient to result in L3 larvae with a clear seizure phenotype (unpaired t-test, *t*(28) = *5.45*, *p < 0.0001, n = 15*, Fig. 2F). These results, in combination, suggest that there are two plasticity windows that overlap during embryogenesis when rearing occurs at 25°C: a BL-CP time locked at 17-19 hrs, and an activity-CP governed by development time and sensitive to a range of activity manipulations (in this instance 32°C).

### Circadian mechanism underlying the timing of the BL-CPs

Two observations are individually suggestive of a possible circadian-based mechanism. First, the BL-CPs are ∼24hrs apart and second, like the clock, they are temperature-compensated (23–25). To investigate this further, we first focused on the potential requirement for CRY, a blue light sensitive clock protein (26). Thus, *cry* null mutant (*cry^02^*) embryos were exposed to BL during either the embryonic or L1 BL-CP. We found that *cry* null embryos that were exposed to BL between 17- 19 hrs (one-way ANOVA, *f*(2, 42) = *1.61*, *p = 0.97, n = 15*) or during L1 at 20-22 hrs ALH did not exhibit a characteristic seizure phenotype at L3 (one-way ANOVA, *f*(2, 42) = *1.87*, *p = 0.93, n = 15*, Fig 3A-B). The involvement of CRY implicates the circadian clock. To validate this, we tested a *period* null mutant (*per^01^*). We found that the BL exposure, during the same two time windows, also did not induce a seizure phenotype (17-19hrs one-way ANOVA, *f*(2, 42) = *10.44*, *p = 0.82, n = 15*), (20-22 hrs ALH one-way ANOVA, *f*(2, 42) = *3.87*, *p = 0.98, n = 15,* Fig. 3C-D). In contrast to this, we found that exposure to 32°C, in *cry^02^*, during 17-19 hrs AEL was sufficient to induce a clear seizure phenotype at L3 (one-way ANOVA, *f*(2, 42) = *2.58*, *p < 0.0001, n = 15*). Interestingly, exposure to 32°C in *per^01^* during 17-19 hrs AEL was lethal.

**Figure 3.**
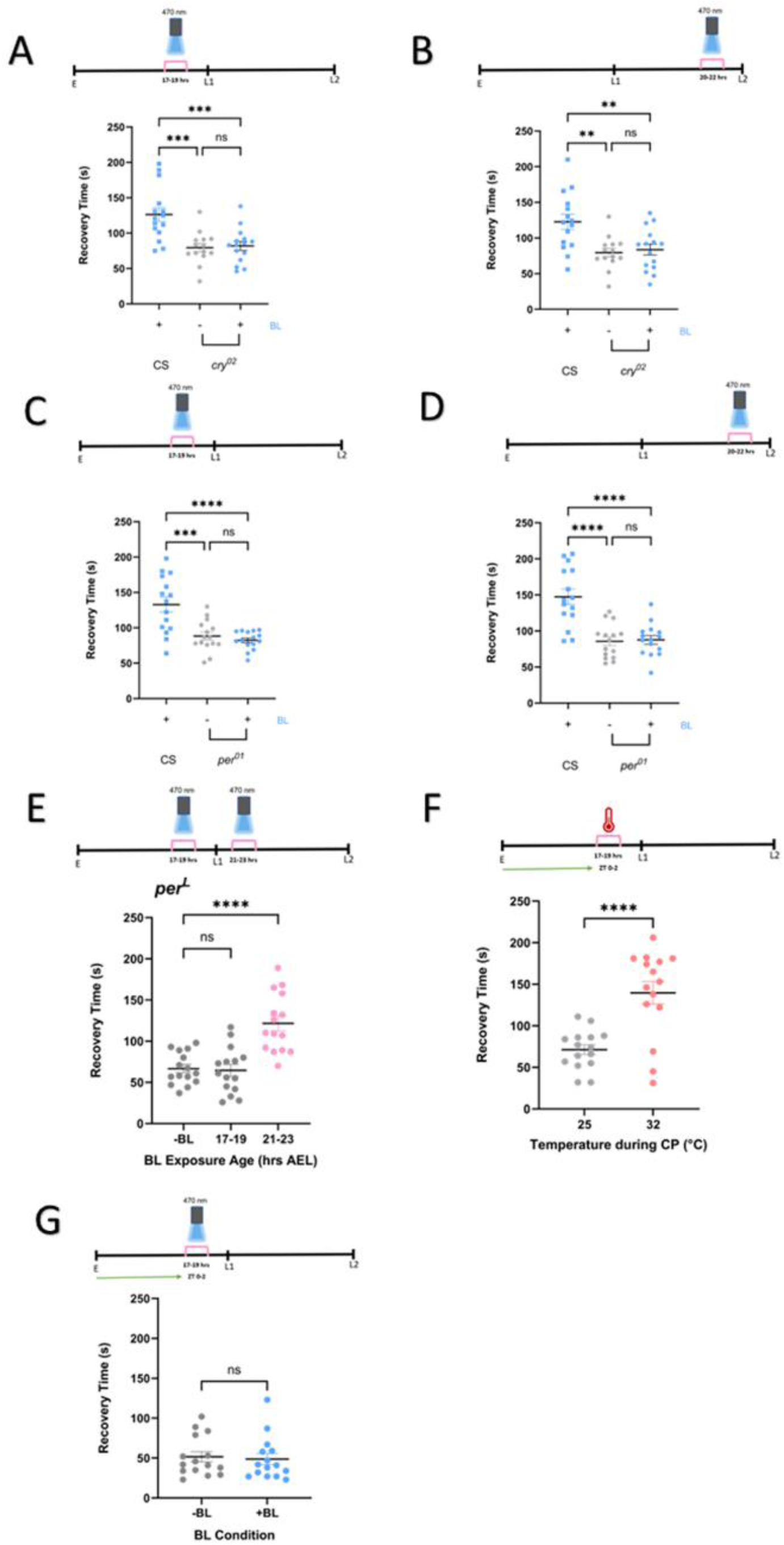
The circadian clock regulates BL-sensitivity. (A) cry02 embryos exposed to BL between 17-19 hrs AEL did not show an increased seizure recovery time compared to dark reared controls (one-way ANOVA, f(2, 42) = 1.607, p = 0.970, n = 15). (B) cry02 late-stage L1 exposed to BL at 20-22 hrs ALH did not have a higher recovery time compared to dark reared controls (one-way ANOVA, f(2, 42) = 1.871, p = 0.931, n = 15). (C) per01 embryos exposed to BL between 17-19 hrs AEL did not exhibit a higher recovery time compared to dark reared controls (one-way ANOVA, f(2, 42) = 10.44, p = 0.825, n = 15). (D) per01 late-stage L1 exposed to BL at 20-22 hrs ALH did not have a higher recovery time compared to dark reared controls (one-way ANOVA, f(2, 42) = 3.867, p = 0.982, n = 15). (E) perL embryos exposed to BL between 21-23 hrs AEL had a higher recovery time compared to dark reared controls (one-way ANOVA, f(2, 42) = 2.038, p < 0.0001, n = 15). (F) CS embryos exposed to 32°C between 17-19 hrs AEL (ZT 0-2), showed an increased recovery time compared to 25°C reared control (Unpaired t-test, t(28) = 4.57, p < 0.0001, n =15). (G) CS embryos exposed to BL between 17-19 hrs AEL (ZT 0-2), did not have an increased recovery time compared to dark reared controls (Mann-Whitney U, U = 99.5, p = 0.602, n = 15).

We next measured effect of BL exposure in two different *per* alleles: *per^Short^ (per^S^)*, with a 5-hr shorter circadian period of 19 hrs, and *per^Long^ (per^L^)*, with a 4-hr longer period length of 28 hrs (27). Interestingly, *per^S^* exhibited an equivalent increased seizure recovery time when dark reared (control) compared to when exposed to BL at the L1 sensitive period. Whilst this points to a potentially interesting genotype-specific effect on neuronal excitability, further investigation is beyond the scope of the current study. By contrast, we observed that *per^L^* embryos exposed to BL at 21-23 hrs AEL (i.e. 4hrs after the normal embryonic period) exhibited an increased seizure recovery time compared to dark reared control (one-way ANOVA, *f*(2, 42) = *2.038*, *p < 0.0001, n = 15,* Fig 3E). Significantly, *per^L^* embryos exposed to BL between 17-19 hrs AEL did not show a seizure phenotype (p = 0.95), indicative of a failure to correctly time the opening of the BL-CP.

To take this investigation one step further, we investigated the potential impact of the ZT exposure time. Thus, we altered the ZT time at which 17-19 hrs AEL occurred. Throughout this study, 17-19 hrs AEL occurs at ZT 18-20, due to laying occurring in the morning. For this specific experiment we adjusted our embryo collection times so that 17-19 hrs AEL occurred at ZT 0-2. This allowed us to separate the potential effect of circadian timing from developmental timing, as 17-19 hrs AEL now occurred at lights on, rather than in the middle of the night when low light is anticipated. First, we confirmed that exposure to 32°C during 17-19 hrs AEL (ZT 0-2) still caused a seizure phenotype (unpaired t-test, *t*(*28*) *= 4.57, p < 0.0001, n =15,* Fig. 3F). This demonstrated that the change in the ZT did not influence the characterized embryonic activity-CP. By contrast, we found that BL exposure during the same ZT-shifted period had no effect on seizure recovery time (Mann-Whitney U, *U = 99.5*, *p = 0.60, n = 15,* Fig 3G). A potential explanation for this change may, conceivably, be the clock’s anticipation of light at ZT 0, demonstrating that the clock is an underlying factor in the BL exposure response. Taken together, our data greatly supports the premise that the BL-CPs have a clear link to, and are possibly regulated by, the circadian clock.

### Clock mediated *cry* mRNA abundance coincides with BL-CPs

To further link the BL-CPs to the circadian clock, we measured mRNA levels for selected clock proteins (*cry*, *tim* and *per*) during both embryonic and larval development. All mRNA expression levels were analyzed using the 2^-ΔΔCt^ method with the first collection, at 12 hrs AEL, being normalized to 1 and used as the reference point. The mRNA expression of *cry* was observed to cycle throughout development, with significant peaks at 18 hrs AEL, 24 hrs ALH, and at 48 hrs ALH (all equivalent to ZT 19). Further larval stages were not examined. We also observed a significant decrease in expression at 0, 12 and 36 hrs ALH. This is clearly indicative of cyclical expression on an ∼24-hour cycle. Importantly, the peaks in *cry* expression coincide with the timing of the BL-CPs that we identify (Fig 4A). By comparison, both *per* and *tim* mRNA levels were found to decrease significantly after 12 hrs AEL and remain low until late L2, thereafter increasing back to initial levels (Fig 4B-C). This suggests that BL-CPs may not require cycling of all clock proteins. However, the clear cycling of *cry* highlights a potential key role for this blue light-sensitive protein.

**Figure 4.**
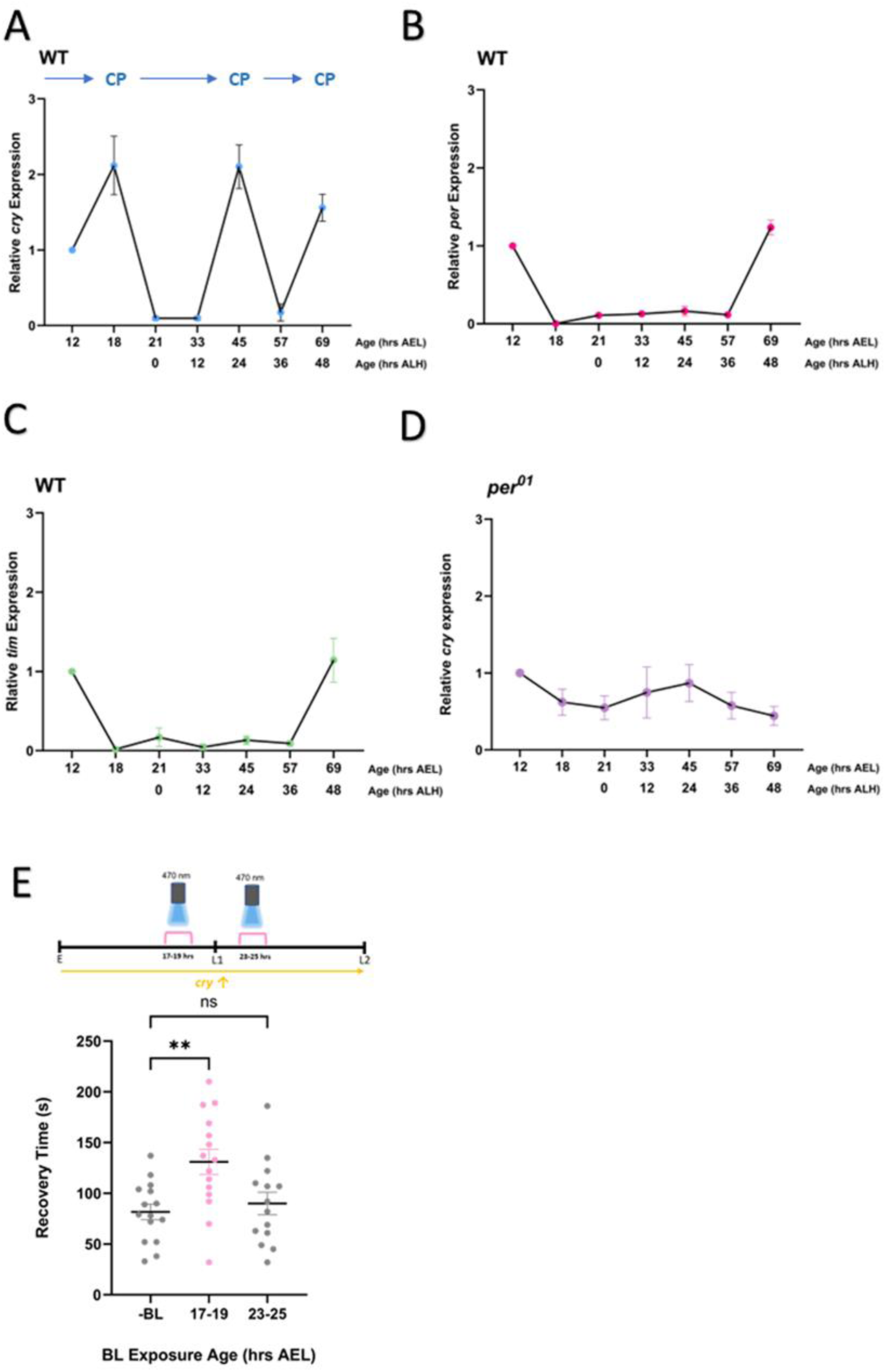
Cry expression coincides with BL-CPs. (A) cry mRNA levels cycle throughout development, with peaks at 18 hrs AEL, 21 hrs ALH and 45 hrs ALH, which coincide with the opening of BL-CPs. (B) per mRNA levels reduce after 12 hrs AEL (set at 1) and remain low until late stage L2, without cycling. (C) tim mRNA levels also reduce after 12 hrs AEL levels and remain low until late stage L2, without cycling. (D) cry mRNA levels did not cycle in a per null mutant. (E) Embryos overexpressing cry and exposed to BL between 17-19 hrs AEL show an increased recovery time at L3, compared to the dark reared control (one-way ANOVA, f(2, 42) = 1.389, p = 0.0035, n = 15). Exposure to BL either side of 17-19 hrs AEL did not, however, induce an increased recovery time (p = 0.806).

A lack of cycling in *per*, yet its necessity for BL-effect is interesting. Possibilities include a requirement of Per to allow *cry* to cycle. Thus, we measured *cry* mRNA expression in the *per^01^* mutant and observed just that, a lack of *cry* cycling in this genetic background (Fig 4D). To determine whether high *cry* expression levels alone are sufficient to support BL-CPs, we pan-neuronally overexpressed *cry* throughout embryonic and larval development (elav-Gal4;;*cry^03^*>w-;UAS-DmCRY;*cry^01^*).

We specifically used a *cry* null background to eliminate endogenous *cry* cycling. We found that overexpression of transgenic *cry* throughout development was not sufficient to open additional CPs (Fig. 4E). Embryos exposed to BL at 17-19 hrs AEL showed an expected increased seizure recovery time (one-way ANOVA, *f*(2, 42) = *1.389*, *p = 0.0035, n = 15*). However, embryos exposed to BL outside of this timing, whilst *cry* expression was equally high, did not (p = 0.806).

### PDF highlighted as a potential signaling pathway

Our data support CRY being essential for BL-sensitivity. This would suggest that CRY-expressing neurons are integral to the signaling pathway underlying the induced seizure phenotype. Because CRY is only expressed in few neurons during larval development (28), there is an assumption that these neurons are coordinating a wider network wide effect. To show this, we inhibited activity of CRY-positive neurons by expressing an inwardly-rectifying potassium channel (Cry39-Gal4>UAS-Kir) (29), and then exposed embryos to BL at 17-19 hrs AEL. Inhibiting activity of CRY neurons was sufficient to prevent BL-induction of a seizure phenotype (one way ANOVA, *f*(2, 42) = *2.887*, *p = 0.25, n = 15*, Fig. 5A).

**Figure 5.**
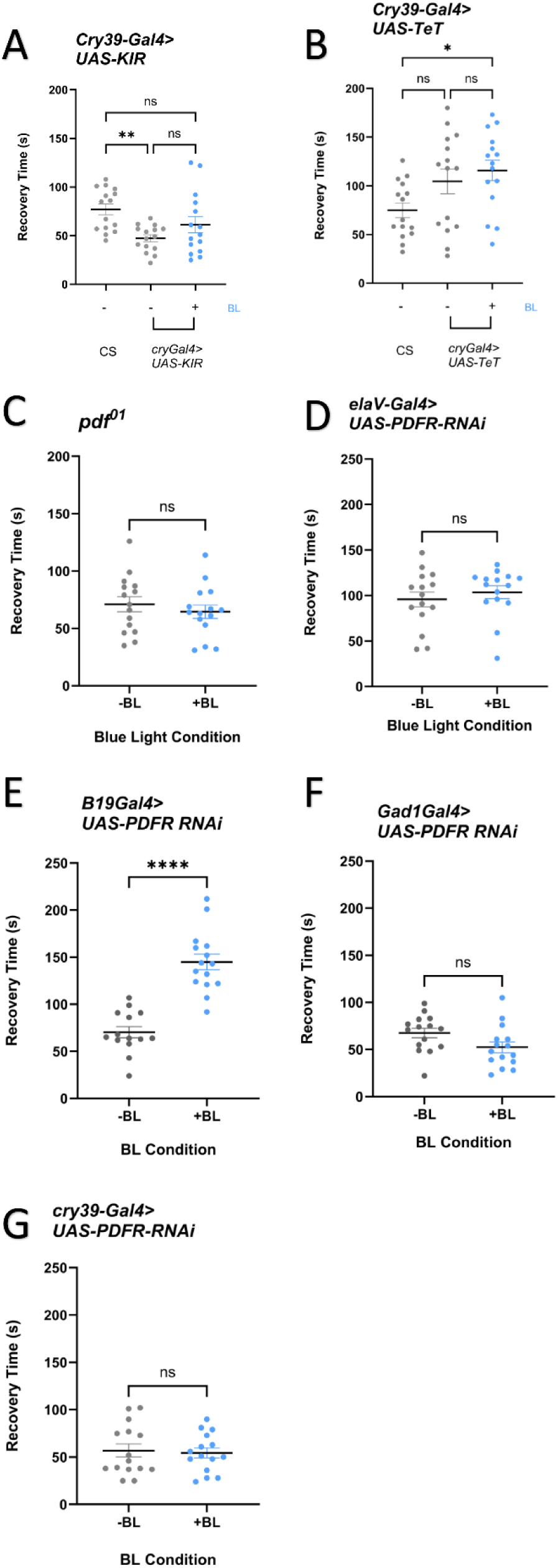
PDF is required for transduction of effect of BL exposure. (A) Silencing activity in CRY-positive neurons, using Kir2.1, is sufficient to prevent induction of a seizure phenotype following BL exposure between 17-19 hrs AEL (one way ANOVA, f(2, 42) = 2.887, p = 0.256, n = 15) (B) Expression of tetanus (TeT) in CRY-positive neurons did not prevent induction of a seizure phenotype following exposure to BL between 17-19 hrs AEL (unpaired t-test, t(28) = 1.577, p = 0.024, n = 15,). However, the presence of TeT alone was sufficient to induce a seizure phenotype in dark-reared animals. (C) pdf01 embryos exposed to BL at 17-19 hrs AEL did not have an increased recovery time compared to dark reared controls (unpaired t-test, t(28) = 0.472, p = 0.472, n = 15). (D) Pan-neuronal PDFR knockdown was sufficient to prevent induction of a seizure phenotype (unpaired t-test, t(28) = 0.738, p = 0.467, n = 15). (E) PDFR knockdown in cholinergic neurons did not prevent induction of a seizure phenotype following exposure to BL at 17-19 hrs AEL (unpaired t-test, t(28) = 7.15, p < 0.0001, n = 15). (F) PDFR knockdown in GABAergic neurons was sufficient to prevent a seizure induction in embryos exposed to BL at 17-19 hrs AEL (unpaired t-test, t(28) = 1.97, p = 0.06, n = 15). (G) PDFR knockdown in CRY positive neurons is sufficient to prevent induction of a seizure phenotype (unpaired t-test, t(28) = 0.299, p = 0.767, n = 15).

It is important to establish how CRY-expressing neurons transduce the effect of BL-stimulation. Thus, we expressed tetanus toxin (Cry39-Gal4>UAS-TeT, (30)) to abolish classical synaptic transmission in CRY neurons. We found that suppressing classical synaptic transmission in CRY neurons was not sufficient to prevent the seizure phenotype induced by BL exposure during the embryonic CP (one way ANOVA, *f*(2, 42) = *1.577*, *p = 0.024, n = 15*, Fig. 5B). Our results are confounded, however, by the fact that expression of tetanus was alone sufficient (i.e. without BL stimulation) to induce a seizure phenotype. Thus, this result remains inconclusive. Studies have shown that tetanus is not always effective to prevent neuronal release of neuropeptides (31). CRY-containing neurons express pigment dispersing factor (PDF, (28)). We used a *pdf* null mutant (*pdf^01^*) and exposed embryos to BL during the 17-19 hrs AEL. The *pdf^01^* embryos exposed to BL did not show an expected increase in seizure recovery time compared to dark reared controls (unpaired t-test, *t*(28) = *0.729*, *p = 0.47, n = 15,* Fig. 5C). Taken together, these observations are consistent with CRY neurons, through the balanced release of both a classical neurotransmitter(s) and a neuropeptide(s), being required to stabilize neural circuit functionality.

Having established a requirement for PDF, we initiated a limited screen to identify target neurons. To do this, we first knocked down PDF receptor expression pan-neuronally using elav-Gal4>UAS-PDFR-RNAi. We found that BL exposure at 17-19 hrs AEL, under these conditions, did not induce a seizure phenotype (unpaired t-test, *t*(28) = *0.738*, *p = 0.47, n = 15,* Fig. 5D). To localize PDF-target neurons, we knocked down PDFRs in excitatory (cholinergic B19-Gal4) or inhibitory (GABAergic, gad1-T2A-Gal4; TM6b) neurons. BL exposure, at 17-19 hrs AEL, in embryos where PDF receptors were knocked down in cholinergic neurons, was still sufficient to increase seizure recovery time compared to dark reared controls (unpaired t-test, *t*(28) = *7.15*, *p < 0.0001, n = 15,* Fig. 5E). By contrast, PDFR knockdown in GABAergic neurons prevented the induced seizure phenotype (unpaired t-test, *t*(28) = *1.97*, *p = 0.06, n = 15,* Fig. 5F).

PDF-containing neurons partake in autocrine signaling (28). We investigated the impact of knocking down PDF-receptors in CRY positive neurons (Cry39-Gal4>UAS-PDFR-RNAi), to assess whether autocrine signaling is integral to the mechanism of this study. Significantly, BL exposure (17-19hrs AEL) in this genotype resulted in no seizure phenotype (unpaired t-test, *t*(28) = *0.299*,*= 0.77, n = 15*, Fig. 5G).

## Discussion

It is established that circadian rhythms regulate most, if not all, aspects of physiology and behavior (15, 32, 33). However, the specific influence of the biological clock on embryonic development remains less well understood. Moreover, clock-mediated effects in developing model systems that are amenable to study are lacking, due to constraints such as *in utero* development. In this regard, our study is significant in that it clearly identifies clock-controlled CPs at 24 hr intervals, from embryo through to larval stages, in the experimentally tractable *Drosophila* model system. Manipulation of these CPs, by exposure to BL, results in a larval seizure phenotype that is a proxy for an altered excitation-inhibition balance within the developing larval locomotor circuitry (34). Thus, our results show a clear clock-mediated effect to neural network development in an embryonic system. In this regard it is interesting to note that the larval stages of holometabolous insects (e.g. *Drosophila*) have been argued to represent free-living embryos: an advantageous strategy that may favor survival of the species (35). This highlights that this study is not only the first instance of a circadian influence on larval development, but also as a potentially more accessible model for further investigation into circadian influence on developing organisms.

Whilst evidence exists to demonstrate the influence of the maternal biological clock on the developing fetus in mammals (for review see (36)), specific effects on the fetus remain speculative. However, disturbance of the fetal circadian system has been linked to increased risk of cardiovascular and other diseases in later life (37, 38). Our observation of CPs in the development of the embryonic and larval nervous system, that are regulated by the clock, provides convincing evidence for circadian influence on a developing organism. In keeping with this hypothesis, it is notable that clock genes, in mice, have been shown to time the postnatal development of neurons pivotal to CP plasticity (39). This demonstrates that the molecular clock, and the plethora of genes under its control have a role in the timing of CPs, providing support for our results. Interestingly, in the same study, it was demonstrated that the pharmacological enhancement of GABA, via exposure to diazepam, was able to overcome delayed neuron development, due to clock gene manipulation, and restore the timing of CP onset (39). This suggests that clock genes may directly control neuronal activity, which in turn is mediated by neurotransmitter release, an area that is explored in the current study.

Clock-based modulation of neuronal activity is well established, in both mammals and *Drosophila* (40, 41). Our results are consistent with manipulation of CPs mediating effects through alterations in release of neurotransmitters. Thus, we show that hyperpolarization of clock neurons, achieved by expression of Kir, is sufficient to prevent effects of BL exposure. The simplest explanation is that hyperpolarization is sufficient to prevent SNARE-mediated neurotransmitter release (29). However, developmental changes in these neurons, that for example might include altered synaptic targeting, remain a possibility. The paradoxical effect of expression of tetanus in clock neurons, which is alone sufficient to induce a seizure phenotype, is intriguing. Coupled with our demonstration that the neuropeptide PDF is required for BL-mediated perturbation of the CPs, our results are consistent with clock neuron activity being tightly regulated to maintain an appropriate balance in transmitter release, or possibly that neurotransmitter release is appropriately timed. Preventing release of a classical transmitter (most likely GABA, which is a key neurotransmitter in the adult clock (42)) achieved by tetanus expression, is sufficient to destabilize the circuit, a similar effect also resulting from BL-mediated release of PDF. How these respective transmitters mediate their effects, and the identity of downstream targets remains to be determined. Downregulating clock genes in PDF-expressing clock neurons, in developing adult *Drosophila*, is sufficient to prevent these cells from fasciculating to form typical axonal bundles (43). As PDF is one of the main signaling outputs of clock neurons (28), it is perhaps not surprising that we implicate this neuropeptide to be a key mediator of BL-exposure of the CPs we identify. Previous studies have demonstrated that small lateral ventral neurons (sLNv) first show robust PDF staining (visible by antibody), along with a small number of other cells, from the first few hrs ALH (44). Therefore, it is logical to propose the sLNvs are likely mediating the effects we observe.

In summary, we identify a series of CPs, regulated by the molecular clock, the manipulation of which by BL significantly impacts neuronal development in the embryo and larva of *Drosophila*. Whilst there is evidence to link the molecular clock with neuronal development in other species and in adult *Drosophila*, this report is the first example in *Drosophila* larvae. This is of significance because this discovery might offer a tractable animal model with which to investigate the role of the molecular clock and circadian activity in the development of an embryonic CNS; something that remains difficult to achieve in the experimentally inaccessible mammalian fetus.

## Materials and Methods

### Fly Stocks

Flies were reared on standard corn meal medium at 25°C under a 12hr-12hr light-dark cycle except for when embryos/larvae were kept in darkness before BL or 32°C exposure, after which they were placed back on this light-dark cycle. Embryos were collected at ZT 1-5 and exposed to stimulation at ZT 18-20, except for experimental circumstances in which the change is noted. Fly strains used are listed in Table 2.

**Table 2.**
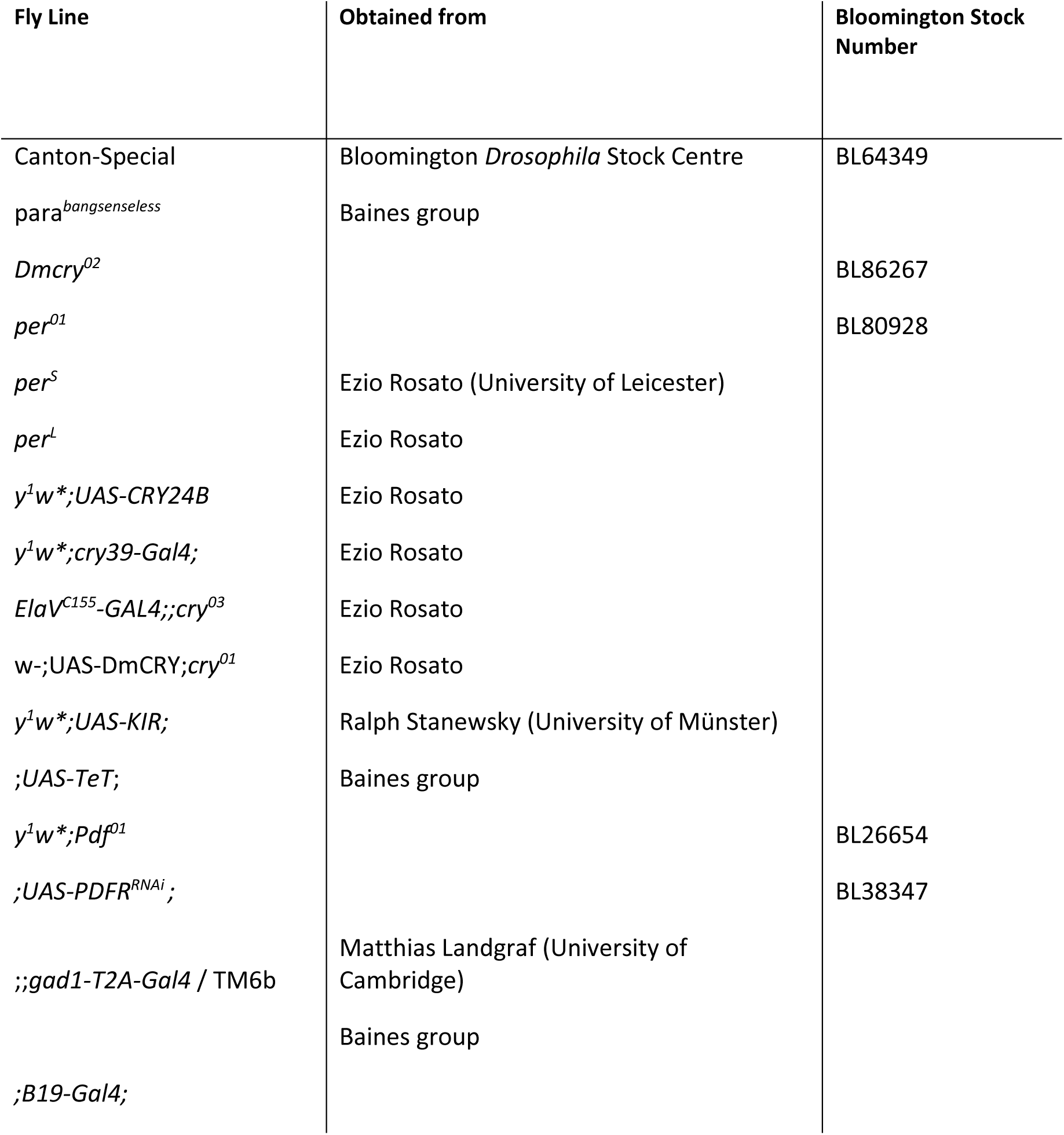
Fly Stock List.

**Table 3.**
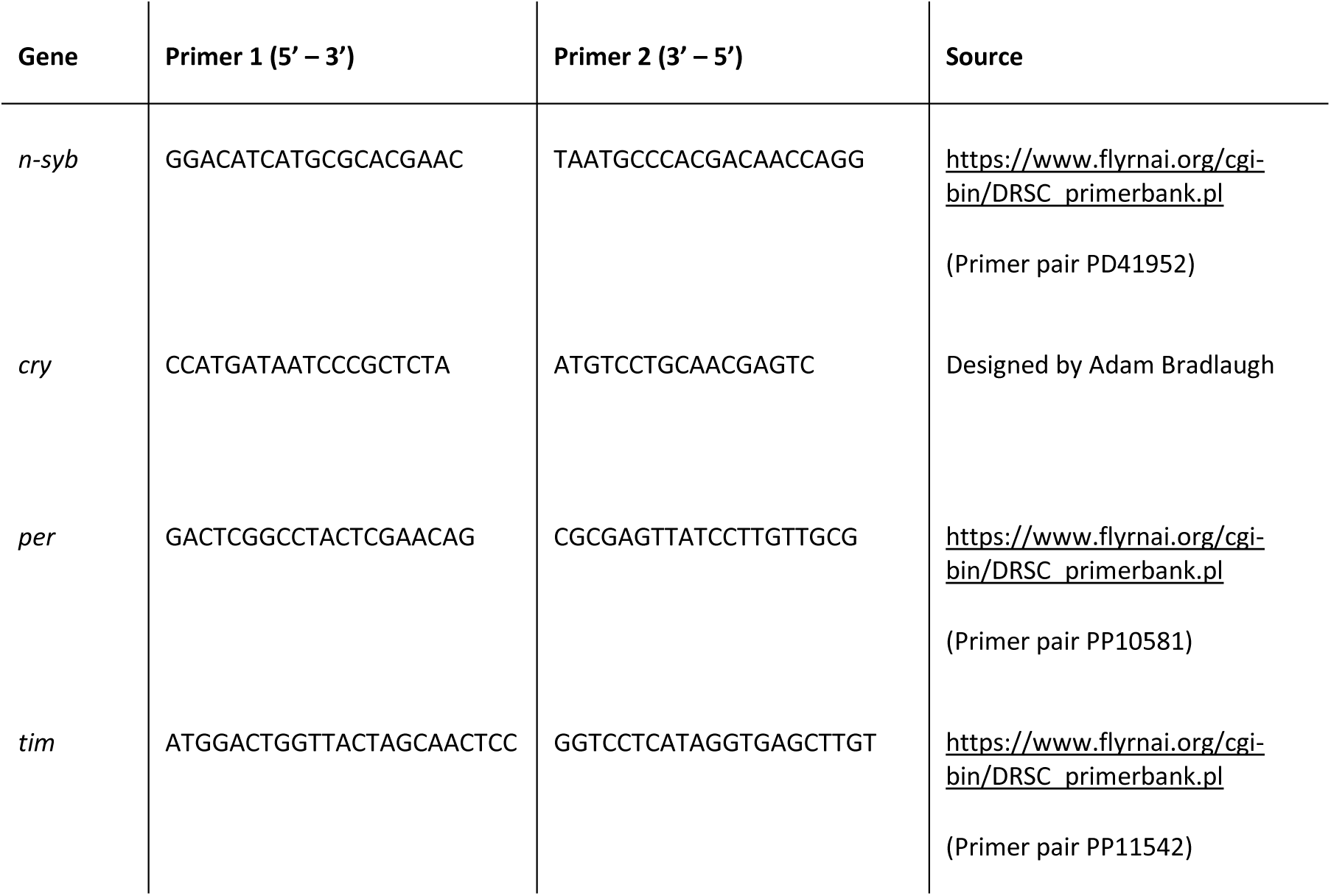
Full primer list.

### Blue light manipulation of neuronal activity

Mated adult females were allowed to lay eggs on grape agar (Dutscher, Essex, UK) plates supplemented with yeast paste at 25°C. Embryos were collected and transferred to a fresh grape agar plate which was subsequently transferred to a 25°C incubator. L1 larvae, when used for manipulation, were collected at 0-1 hours after larval hatching (ALH), by first removing any hatched larvae at lights on (ZT0) and then collecting any newly hatched larvae at ZT1-2. Embryos or larvae were exposed to collimated BL from an overhead LED (8 Wcm^2^). The LED had a peak emission at 470 nm (bandwidth 25 nm, OptoLED, Cairn Instruments, Kent, UK). Light was pulsed for 100msec duration at 2 Hz. Manipulated embryos/larvae were transferred to standard corn meal medium containing vials and reared at 25°C under a 12hr-12hr light-dark cycle until wandering L3.

### Temperature manipulation

Mated adult females were allowed to lay eggs on grape agar plates supplemented with yeast paste at 25°C. Embryos were collected and transferred to an empty PCR tube (0.2ml Eppendorf) which was subsequently placed into a PCR machine (Eppendorf Mastercycler Gradient). When larvae were used in the protocol, these were collected as described above. Embryos/larvae were exposed to 32°C at the appropriate time by programming the PCR machine to warm to this temperature. For embryos exposed to high temperatures at 17-19 hours AEL, the PCR thermal profile was 25°C for 13 hours, 32°C for 6 hours and hold at 25°C. For larvae exposed to high temperatures at 20-22 hours ALH, larvae were placed into PCR tubes at a later stage (6 hours ALH) to minimize lethality due to starvation and desiccation, prior to this they were maintained at 25°C on a grape agar plate supplemented with yeast paste. The larval PCR thermal profile was 25°C for 13 hours, 32°C for 3 hours, hold at 25°C. A small amount of water was added to each sample tube to increase survival rates. Manipulated embryos/larvae were transferred to standard corn meal medium containing vials and reared at 25°C under a 12hr-12hr light-dark cycle until wandering L3.

### Low temperature rescue of neuronal activity

Embryos or larvae were collected and transferred to a fresh grape agar plate as previously described. Embryos were placed in an 18°C incubator for the duration of embryogenesis (determined by the hatching of larvae) or for the duration of L1 (full development at this lower temperature being 44 hours). The embryos/larvae were transferred to standard corn meal medium containing vials and reared at 25°C under a 12hr-12hr light-dark cycle until wandering L3.

### Seizure induction

Wandering L3 larvae were washed in ddH_2_O to remove food residue and subsequently placed onto a clean plastic dish and gently dried with a paper towel. Once normal crawling behavior was observed, a conductive probe composed of two tungsten wires (0.1 mm diameter, ∼1-2 mm apart) was positioned over the approximate position of the CNS. A 3 V DC pulse for 2 s, generated by an isolated constant voltage stimulator (Digitimer Ltd. Uk) was applied. In response to the stimulus, larvae initiated a transitory paralysis with occasional spasms and rolling behavior. The time to resume normal crawling behavior was measured as recovery time (RT). Normal crawling behavior was defined as three consecutive whole body peristaltic waves resulting in forward movement (45). Videos of this phenotype are available in (19).

### RTqPCR

Embryos/larvae were collected as previously described. The embryos were placed into 2 ml Eppendorf tubes and left to mature at 25°C in a normal LD cycle. Once the embryos/larvae had reached the required age, they were flash frozen using liquid nitrogen and stored at -80°C. It is important to note that freezing took place under the appropriate light conditions consistent with the rearing LD cycle. The Qiagen RNeasy Mini Kit was used to extract RNA following the standard protocol (#74004, Qiagen, Germany). RNA was converted to cDNA using the ThermoScientific RevertAid First Strand cDNA Synthesis Kit (#K1622, Thermo Fisher Scientific, UK) following the standard protocol: 10 µl total RNA was incubated with 1µl random hexamers and 1 µl Oligo(dt) at 65°C for 5 minutes then stored on ice. To this mixture 4µl 5x Reaction Buffer, 2µl 10mM dNTP, 1µl RNase inhibitor, and 1µl reverse transcriptase was added. This was incubated at 25°C for 10 minutes, 42°C for 60 minutes, then 70°C for 10 minutes and subsequently stored at - 20°C. The 20µl resultant cDNA product was diluted to 40µl using RNA-free water, prior to RT-qPCR. RT-qPCR was carried out using LightCycler® 480 (Roche Diagnostics, Germany) with 10µl LightCycler® 480 SYBR Green I Master mix (#04707516001) 0.5µl forward primer (5pmol/µl), 0.5 µl reverse primer (5pmol/µl), 1µl single stranded cDNA, and 8µl ddH_2_O in each well of a 96-well plate (#04729692001, Roche). The same thermal profile was used for all primers: 10 minutes at 95°C followed by 55 cycles of [10 seconds at 95°C, 10 seconds at 60°C and 10 seconds at 72°C] (46). All primer pairs used are listed in Table 2. N-syb was used as the reference gene. Data were normalized using the 2^-ΔΔCT^ method, such that the relative expression of each genotype = 1. For each genotype, *n = 3* independent biological replicates, comprising 2-3 technical repeats per replicate.

## Acknowledgments

We thank Ezio Rosato, Ralf Stanewsky and Matthias Landgraf for their generous provision of *Drosophila* strains. We thank Matthew Cobb and Mino Belle and Nara Muraro for comments on a draft of this paper. SD would like to thank Wei-Hsiang Lin (University of Manchester) for training in qRT-PCR. We are also grateful to the members of the Baines group for support and ideas. SD was funded through a BBSRC DTP award provided by the University of Manchester. AAB and RAB were funded through an award from BBSRC to RAB (BB/V005987/1). Work on this project benefited from the Manchester Fly Facility, established through funds from the University and the Wellcome Trust (087742/Z/08/Z).

## Data Availability Statement

Figures show all raw data values in addition to means and sem. Fly stocks used are available on request.

## Notes

### Competing Interest Statement

The authors have declared no competing interest.

### Summary of Updates

We have enlarged the figures to improve readability. We have also revised figure 5B-C as control data was previously incorrect.

